# The anti-inflammatory activity of IgG requires the co-engagement of types I and II Fc receptors

**DOI:** 10.1101/2024.12.16.628704

**Authors:** Andrew T. Jones, Tetyana Martynyuk, Alessandra E. Marino, Stylianos Bournazos, Jeffrey V. Ravetch

## Abstract

Intravenous immunoglobulin administered at high doses has been used for decades as an effective anti-inflammatory preparation to treat a wide array of autoimmune diseases. Studies in murine models have found this activity to be dependent on 1) sialylation of the N-linked glycan on the CH2 domain of IgG, 2) the type I IgG inhibitory Fc receptor FcγRIIB, and 3) the type II Fc receptor DC-SIGN. Here, we demonstrate that the ectodomain glycan on FcγRIIB interacts with the lectin domain of DC-SIGN to form a cell surface complex, thereby enabling the binding of sialylated IgG1. We have exploited this observation to Fc-engineer sialylated IgG1 to enhance its affinity for FcγRIIB and demonstrate that the resulting molecule is at least 100-fold more potent in reducing the inflammatory sequelae of antibody or T cell mediated autoimmune diseases, providing the basis for a new class of anti-inflammatory therapeutics.

## Main Text

Intravenous immunoglobulin (IVIG), a preparation of pooled human IgG collected from thousands of donors, has been a decades-long FDA approved therapeutic for the treatment of a wide variety of autoimmune diseases such as immune thrombocytopenia (ITP), rheumatoid arthritis, chronic inflammatory demyelinating polyneuropathy (CIDP), and Kawasaki’s Disease. It is widely used in over 80 autoimmune diseases (*1*), including neuroinflammatory diseases like myasthenia gravis and blistering diseases like pemphigus (*2*) to reduce the inflammatory sequelae of these diseases. However, the high dose requirement (1-2 g/kg), difficulty in administration, high cost, and availability issues have prompted the need to identify a readily available, recombinantly expressed, replacement product. Studies in animal models have established the requirement of terminal α2,6 linked sialylation of the single N-linked glycan (N297) in the IgG1 Fc-domain in mediating the anti-inflammatory activity of IVIG (*3, 4*). Furthermore, genetic knockout studies in murine models have implicated both type I (canonical, IgG superfamily members), and type II (C-type lectin) IgG Fc receptors (FcγRs) in the mechanism by which sialylated IgG mediates its activity *in vivo* (*3, 5-7*). Recombinantly expressed sialylated IgG1 Fc (sFc) or an Fc mutant that phenocopies the structural perturbation of the sialylated IgG1 CH2 domain (F241A Fc) (*8, 9*), show *in vivo* potency at 100 mg/kg (*10*) doses, a 10-20-fold enhancement in effective dose compared to IVIG in the same models.

Most studies characterizing IVIG *in vivo* utilize murine models expressing endogenous murine FcγRs, which have distinct functions, expression patterns, and binding affinities for IgG1, compared to human FcγRs (*11*). Here, we characterize the anti-inflammatory activity of IVIG and sialylated IgG in FcγR humanized mice, which recapitulate the expression, function, and binding patterns of human FcγRs on the murine background (*12*), and determine the contributions of type I and type II FcγR engagement and signaling to the *in vivo* potency of IVIG. These studies result in a novel sialylated Fc variant with at least 100-fold enhanced potency, compared to IVIG, and describe a novel function of type II FcγRs, which is to directly associate with type I FcγRs and mediate their binding to sialylated IgG.

### Requirements of sialylation and type I FcγR engagement in hFcγR mouse models of IVIG mediated anti-inflammatory activity

We first determined if IVIG and recombinant sialylated wild-type IgG1-Fc (WT sFc) mediate anti-inflammatory activity in hFcγR mice, and if this activity is sialylation dependent. To generate recombinant WT sFc proteins, we co-transfected HEK 293-F cells with WT IgG1 Fc expression vectors and plasmids encoding the genes for Beta-1,4-galactosyltransferase 1 (B4GALT1) and Beta-galactoside alpha-2,6-sialyltransferase 1 (ST6GAL1), which attach galactose and α2-6 linked sialic acid, respectively, to the terminal N-Acetylglucosamine (GlcNAc) on the complex, biantennary N297 glycan (**Fig. 1A)**. This method of co-expressing B4GALT1 and ST6GAL1 with antibody or Fc expression vectors has been shown to result in an 80-90% sialylated product (*9, 13*). We validated the glycan profile of this product by probing with lectins specific for terminal galactose (ECL, Erythrina Cristagalli Lectin) or α2-6 linked sialic acid (SNA, Sambucus Nigra Lectin) (**Fig. 1B, fig. S1A-B)** (*14*). Treating the resulting sFc with neuraminidase cleaves sialic acid from the glycan, resulting in a terminally galactosylated but non-sialylated Fc.

**Figure 1:**
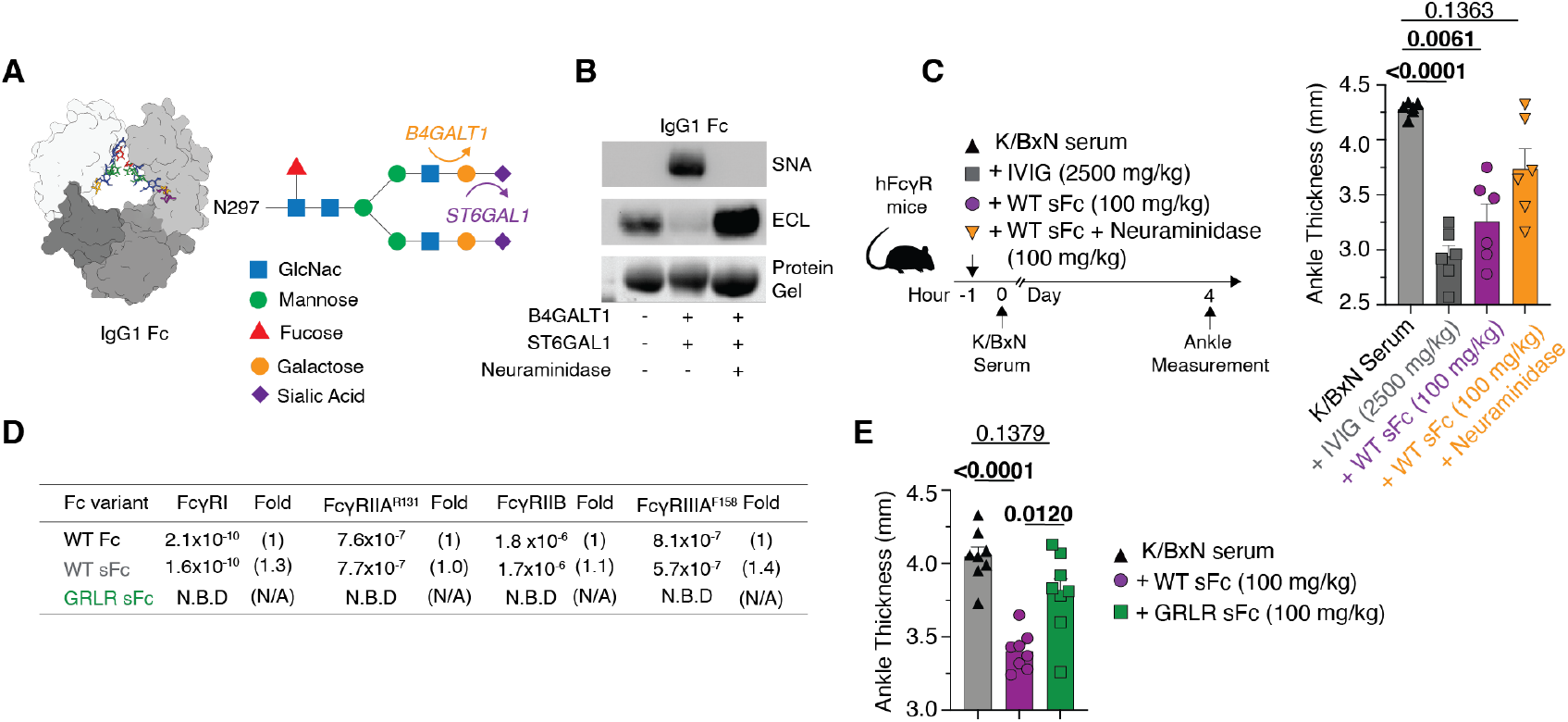
Type I Fc receptor engagement is required for the anti-inflammatory activity of sialylated IgG. **(A)** Structure of the fully processed Asn297 glycan in the human IgG1 antibody Fc domain. The sugars galactose and α2,6 sialic acid are linked to the core glycan via the glycotransferases Beta-1,4-galactosyltransferase 1 (*B4GALT1*) and Beta-galactoside alpha-2,6-sialyltransferase 1 (*ST6GAL1*), respectively. **(B)** Lectin blot analysis of recombinantly produced hIgG1-WT Fc proteins, expressed by itself or along with B4GALT1 and ST6GAL1, as well as neuraminidase treated sFc. Terminal a2,6 sialic acid and galactose detected by the Sambucus Nigra (SNA) lectin and Erythrina Cristagalli (ECL) lectin, respectively. **(C).** hFcγR mice (males, 3/group) were dosed with IVIG (25000 mg/kg), WT sFc (100 mg/kg) or neuraminidase treated sFc (100 mg/kg) one hour prior to injection with K/BxN serum. Right, ankle measurements (Mean +/-SEM) four days post K/BxN serum transfer. **(D)** Table of binding affinities (KD) of WT Fc WT sFc and GRLR sFc for type I FcγRs, determined by SPR. Fold change compared to WT Fc indicated. N.B.D, no binding detected. **(E)** Ankle measurements (mean +/-SEM) on day four of hFcγR mice (male, 4/group) dosed with 100 mg/kg of WT sFc or GRLR sFc one hour prior to K/BxN serum injection. Brown-Forsythe and Welch ANOVA tests, followed by Dunnett’s T3 multiple comparisons test was performed for (**C)** and **(E)**. P-values indicated above groups, significant values in bold.

To characterize the *in vivo* anti-inflammatory activity of IVIG or recombinantly generated sFc in hFcγR mice, we used the K/BxN serum transfer induced arthritis (STIA) model. K/BxN mice spontaneously develop arthritis and swollen joints, and the transfer of serum derived from these mice to naive hosts results the development of transient arthritis which can be evaluated through clinical scoring, histological evaluation of the joint, and measurement of ankle thickness due to inflammation (*15*). Previous studies have demonstrated that high-dose IVIG (1-2.5 g/kg) suppresses K/BxN serum-mediated inflammation in a sialylation-dependent manner, and this activity can be phenocopied using recombinantly produced WT sFc at a 10-30 fold lower dose compared to IVIG (*3, 4, 6*).

FcγR humanized mice were dosed with IVIG (2.5 g/kg), WT sFc (100 mg/kg), or neuraminidase treated WT sFc (100 mg/kg), prepared as described above, one hour before administration of K/BxN serum, and ankle thickness was measured four days later at the time of peak disease. Prophylactic IVIG treatment protected hFcγR mice from developing arthritis, and this effect was phenocopied with WT sFc at a 25-fold lower dose (100 mg/kg) (**Fig. 1C, fig. S1C)**. Notably, neuraminidase treated sFc was unable to protect mice from inflammation at an equivalent dose, demonstrating that sialylation of the Fc glycan is required for anti-inflammatory activity in hFcγR mice in this model.

We next determined if type I FcγR engagement by sFc is required to drive anti-inflammatory activity in hFcγR mice. A common method of knocking out type I FcγR binding to IgG1 Fc is by cleaving the hIgG1-N297 glycan, either through mutagenesis or treatment with glycosidases, as this glycan is required for FcγR binding (*16, 17*). However, this would not allow us to characterize sialylated IgG in the absence of type I FcγR binding. Instead, we used a well-studied pair of mutations (G236R/L328R, GRLR), which ablates type I FcγR engagement, leaves the N297 glycan intact, and has the same half-life and thermostability as WT IgG1 (*18-20*) (**fig. S2A-B)**. We generated sialylated hIgG1-GRLR Fc (GRLR sFc), characterized its sialylation status by western blotting with SNA and ECL (**fig. S2C-D)**, and confirmed its lack of binding to type I FcγRs by surface plasmon resonance (SPR) (**Fig. 1D, fig. S2E)**. Next, we dosed hFcγR mice with 100 mg/kg of WT sFc or GRLR sFc one hour prior to administration of K/BxN serum and determined disease severity four days later. Unlike WT sFc treated mice, GRLR sFc treatment failed to protect mice from arthritis (**Fig. 1E)**, indicating that type I FcγR engagement, along with sialylation, are both required to mediate the anti-inflammatory activity of IgG in FcγR humanized mice.

Sialylation of mouse IgG subclasses, notably mouse IgG1 and mouse IgG2b, has been shown to decrease their affinities for activating mouse FcγRs while maintaining native affinities for the inhibitory receptor FcγRIIB, resulting in a shift towards engagement of inhibitory over activating FcγRs(*3*). We performed SPR to measure binding affinities of human FcγRs to Protein G immobilized sFc or neuraminidase treated sFc. Unlike mouse IgG, sialylation of human IgG1 Fc did not significantly impact its binding to type I human FcγRs, and thus cannot account for the enhanced anti-inflammatory activity of sFc compared to neuraminidase treated sFc (**fig. S3A-D)**, consistent with a role for other receptors, such as the Type II FcγRs, in the anti-inflammatory activity of sialylated IgG.

### Targeting FcγRIIB through Fc-engineering enhances the anti-inflammatory activity of sialylated IgG

Canonical type I FcγRs are a group of activating and inhibitory IgG binding receptors expressed on a diverse array of immune cells. Engagement of ITAM-signaling activating FcγR pathways, such as FcγRIA (CD64), FcγRIIA (CD32a), and FcγRIIIA (CD16a) conveys classical Fc-effector functions such as antibody dependent cellular phagocytosis (ADCP), antibody dependent cellular cytotoxicity (ADCC), and innate immune cell activation (*11*). Conversely, the ITIM-signaling FcγRIIB (CD32b) acts as a negative regulator of the immune response, safe-guarding against hyper-immune activation. FcγRIIB is required for IVIG to mediate anti-inflammatory activity *in vivo*, as mice in which FcγRIIB has been blocked or genetically knocked out are unresponsive to IVIG therapy in multiple models of autoimmunity (*5, 21*). Most innate immune cells expressing activating FcγRs co-express FcγRIIB, and the balance of signaling between activating and inhibitory receptors determines a threshold for cell activation (*22*). Furthermore, we have described a mechanism in which infusion of IVIG or sFc induces the up-regulation of FcγRIIB on innate immune cells, raising the threshold for FcγR-mediated pro-inflammatory responses (*6*). This up-regulation of FcγRIIB in response to IVIG therapy has also been observed in patients with CIDP, a disease in which IVIG serves as a first-line therapy (*23*).

While we have proposed FcγRIIB engagement and immunomodulation as a primary mechanism of IVIG mediated anti-inflammatory activity, alternative mechanisms have been proposed which argue IVIG mainly functions by actively blocking activating FcγRs or the neonatal Fc receptor (FcRn), competing with pathogenic autoantibodies for FcγR signaling (*24, 25*) or FcRn mediated antibody recycling (*24*). To determine if targeting inhibitory or activating type I FcγR pathways by sFc mediates anti-inflammatory activity, we generated sialylated versions of two well defined IgG1 Fc variants, the V11 (G237D, P238D, H268D, P271G, A330R) (*26*) and GA (G236A) (*20*) Fc proteins which have enhanced affinity for inhibitory or activating type I FcγRs, respectively (**fig. S4A-D**). Compared to WT sFc, GA sFc has an approximately 10 fold enhanced affinity for the activating FcγRIIA while having comparable affinities for FcγRIIB and FcγRIIIA, while the V11 sFc has a 37-fold enhancement in affinity for the inhibitory FcγRIIB **(Fig. 2A)**. Additionally, V11 sFc binds to the high affinity FcγRIA with 193-fold weaker affinity compared to WT sFc, and is completely unable to bind the H131 variant of FcγRIIA as well as both variants (F158 and V158) of FcγRIIIA (**fig. S5)** (*26*).

**Figure 2:**
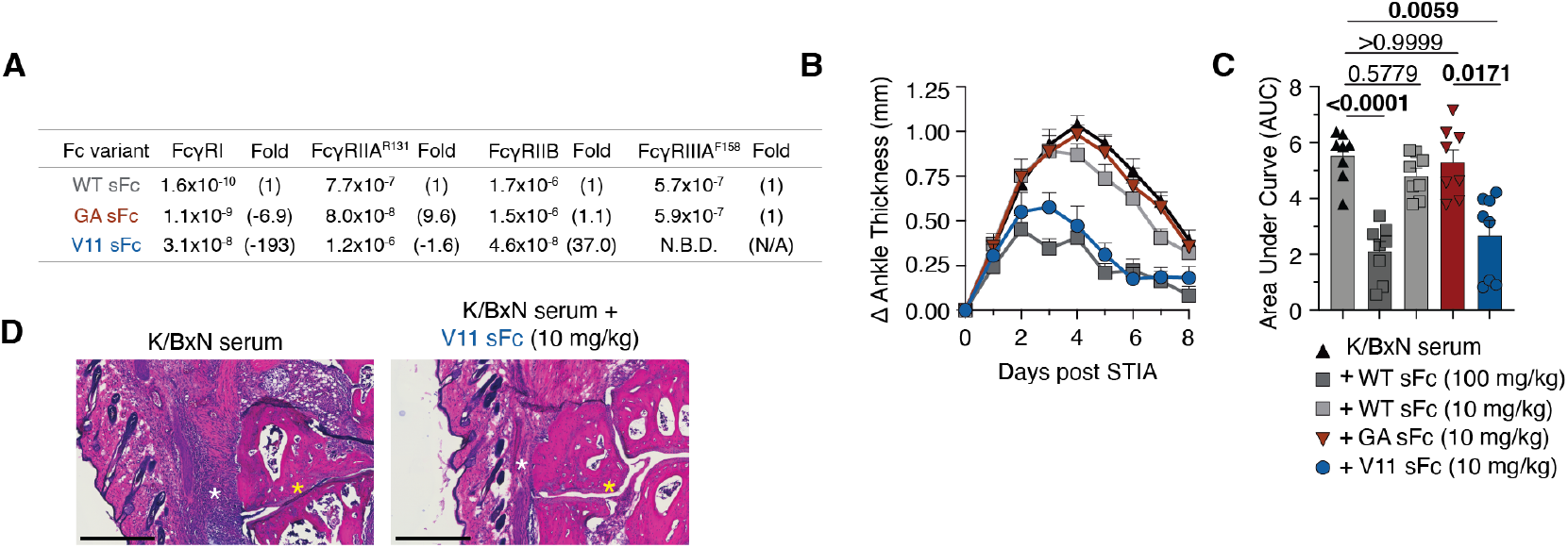
Fc-engineering IgG to target FcγRIIB potently enhances the anti-inflammatory activity of sialylated IgG. **(A)** Table of binding affinities (KD) of WT sFc, GA sFc or V11 sFc to type I FcγRs, determined by SPR. Fold change in affinity compared to WT sFc indicated. **(B)** Kinetics of change in ankle thickness in hFcγR mice (4/group) dosed with WT sFc (100 mg/kg), WT sFc (10 mg/kg), GA sFc (10 mg/kg) or V11 sFc (10 mg/kg) one hour prior to K/BxN serum injection. Mice were monitored over the subsequent eight days with daily ankle measurements. **(C)** Area under curve (AUC) analysis of ankle kinetics in each group. Brown-Forsythe and Welch ANOVA tests, followed by Dunnett’s T3 multiple comparisons test. P-values indicated above groups, significant values in bold **(D).** Hematoxylin and Eosin (H&E) staining of ankle joints in mice with K/BxN serum induced arthritis, untreated or treated with V11 sFc (10 mg/kg). White asterisk denotes immune cell infiltration into joints. Yellow asterisk highlights the decreased space between bones in the ankle joint. Scale bar, 500 um.

As both GA and V11 sFc have enhanced affinities for activating or inhibitory FcγRs, respectively, we theorized they may mediate anti-inflammatory activity at a 10-fold lower dose (10 mg/kg) compared to WT sFc. Thus, we dosed hFcγR mice with 100 mg/kg or 10 mg/kg of WT sFc, 10 mg/kg of GA sFc, or 10 mg/kg of V11 sFc before administration of K/BxN serum. Mice were then monitored and scored over the next eight days to characterize onset, peak, and resolution of inflammation. Treating mice with 10 mg/kg of V11 sFc mediated potent anti-inflammatory activity, comparable to 10-fold higher dose of WT sFc, suggesting that targeting the inhibitory FcγRIIB pathway drives anti-inflammatory activity **(Fig. 2B-C)**. Histological analysis of ankle joints in V11 sFc treated mice show a marked reduction in immune cell infiltration and increased joint space compared to untreated mice (**Fig. 2D)**. Unlike V11 sFc, GA sFc was ineffective in controlling inflammation, suggesting that targeting or blocking activating FcγR pathways is not responsible for the anti-inflammatory activity of IVIG or sialylated IgG. Furthermore, the lack of binding of V11 sFc to the activating FcγRIIIA suggests that targeting or blocking this receptor is unnecessary to drive anti-inflammatory activity as well. Interestingly, the anti-inflammatory activity of V11 sFc was found to be partially independent of sialylation, as neuraminidase treated V11 sFc failed to protect mice from controlling the early onset of K/BxN serum induced inflammation, measured three days post K/BxN serum injection (**fig. S6A)** but effectively suppressed inflammation at the day six timepoint **(fig. S6B)**. The increased affinity for FcγRIIB may account for the sialylation independent activity of the V11 Fc, though it is notable that some mutations in the V11 Fc (G237D, P238D) are located in a similar region as the F241A Fc variant, which structurally phenocopies sFc and mediates anti-inflammatory activity in a sialylation independent manner (*27*).

We next determined if V11 sFc retains activity at lower doses than 10 mg/kg. We performed a titration study in which we dosed hFcγR mice with 10 mg/kg, 5 mg/kg, or 1 mg/kg of V11 sFc one hour before K/BxN serum injection. As a comparison, an additional group of mice was dosed with a standard therapeutic dose (1000 mg/kg) of IVIG. Treating mice with 10 mg/kg of V11 sFc again provided potent anti-inflammatory activity, blunting both the magnitude and duration of inflammation, and phenocopying the effects of IVIG at a 100-fold lower dose **(Fig. 3A-B)**. However, we found that this activity is lost when V11 sFc is dosed at 1 mg/kg, though treatment with 5 mg/kg of V11 sFc did result in a modest improvement to the resolution of inflammation. These studies demonstrate the effective dose of V11 sFc to be at least 10 mg/kg in FcγR humanized mice, a 100-fold lower dose compared to IVIG. This highly translatable dose is comparable to recent FDA approved engineered IgG1-Fc based therapeutics, which function as a blockade of the neonatal Fc receptor (FcRn) mediated antibody recycling pathway, resulting in the transient depletion of both pathogenic and non-pathogenic antibodies in patients (*9*).

**Figure 3:**
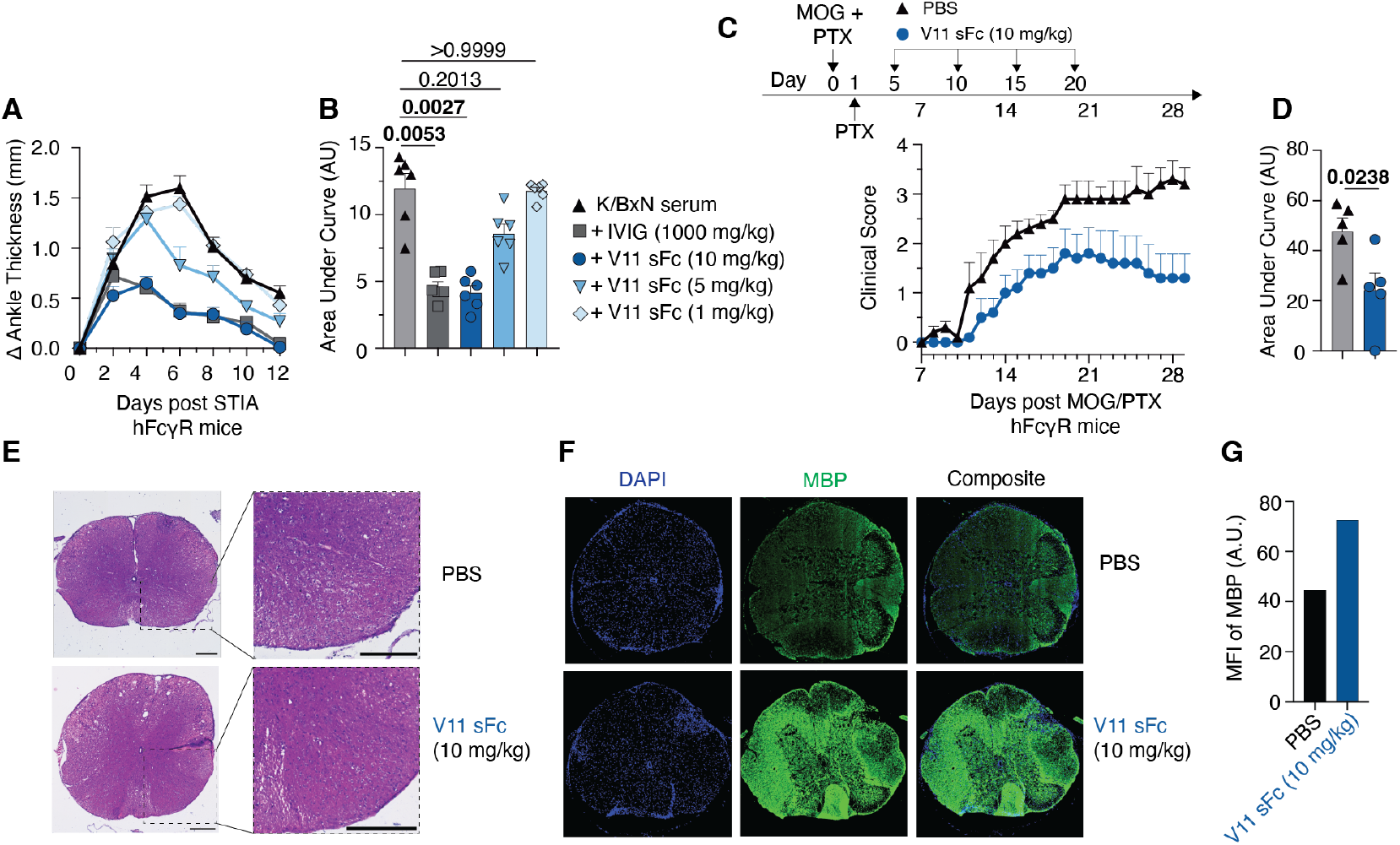
V11 sFc phenocopies the anti-inflammatory activity in at least 100-fold lower doses and ameliorates neurological autoimmunity in a mouse model of multiple sclerosis. **(A).** Kinetics of change in ankle thickness in hFcγR mice (3/group) dosed with IVIG sFc (1000 mg/kg), V11 sFc (10 mg/kg), V11 sFc (5 mg/kg) or V11 sFc (1 mg/kg) prior to K/BxN serum injection. Mice were monitored over the subsequent twelve days with daily ankle measurements. **(B)** Area under curve (AUC) analysis of ankle kinetics in each group. Brown-Forsythe and Welch ANOVA tests, followed by Dunnett’s T3 multiple comparisons test. P-values indicated above groups, significant values in bold. **(C)** hFcγR mice (5/group) were immunized with MOG_35-55_ peptides emulsified in CFA on day 0, along with two subcutaneous injections of pertussis toxin (PTX) on days 0 and 1. Mice were then injected with PBS or V11 sFc (10 mg/kg) on days 5, 10, 15, and 20. Daily clinical scoring to detect neurological symptoms started on day 7, mice were monitored until day 28. **(D)** Area under curve (AUC) analysis of clinical scores in PBS or V11 sFc treated mice in (C). Mann Whitney test. **(E)**. H&E staining of spinal cord sections of untreated and V11 sFc treated mice. Scale bar, 250 um. **(F)** Detection of myelin basic protein (MBP) in spinal cord sections in untreated and V11 sFc treated mice, determined via immunofluorescence. Nuclei staining (DAPI) in blue, MBP in green. **(G)** Quantified relative expression of MBP in (F).

As IVIG is a first line therapy for several neurological autoimmune diseases, such as CIDP, Guillain–Barré syndrome (GBS), and multifocal motor neuropathy (MMN), we next determined in V11 sFc could control inflammation in mouse models of autoimmune neuropathy (*28*). IVIG has been demonstrated to protect mice from developing neurological inflammation in the experimental autoimmune encephalomyelitis (EAE) model of multiple sclerosis, primarily through promoting the expansion of T-regulatory cells (*8, 29, 30*). To determine if V11 sFc protects mice from neuro-inflammation, we induced EAE by immunizing hFcγR mice with a myelin oligodendrocyte glycoprotein (MOG) derived peptide MOG_35-55_ emulsified in complete Freund’s adjuvant (CFA) along with pertussis toxin (PTX). Mice then received four doses of 10 mg/kg of V11 sFc, starting on day five, spaced five days apart. Treatment with 10 mg/kg V11 sFc significantly lowered clinical scores in mice, compared to untreated controls, suggesting that V11 sFc is effective in controlling neuro-inflammation and demyelination (**Fig. 3C-D)**. Additionally, we performed immunofluorescence and histochemical analysis on spinal cord sections of untreated and V11 sFc treated mice to determine the degree of demyelination, measured by immunofluorescent based detection of myelin-basic protein (MBP), as well as inflammation and cell destruction, measured by H&E staining. We found V11 sFc treatment prevented cell destruction in spinal cords (**Fig. 3E)**, and retained high levels of MBP staining, which was significantly reduced in untreated mice **(Fig. 3F-G)**. The ability of V11 sFc to mediate anti-inflammatory activity in multiple models of both passively transferred and endogenously induced autoimmune diseases highlights its potential as a potent anti-inflammatory therapeutic with broad applications.

### Type I and II FcγRs synergize to bind IgG and drive anti-inflammatory activity

A distinct class of Fc receptors, type II FcγRs are C-type lectin related receptors found on diverse subsets of immune cells including B-cells and myeloid-lineage derived innate immune cells, all of which co-express type I FcγRs (*11*). The type II FcγR DC-SIGN (CD209), or its mouse homolog SIGN-R1 (CD209b), is required for IVIG and sFc to mediate anti-inflammatory activity, as mice lacking these receptors fail to control inflammation in multiple models of autoimmunity (*6, 9, 31*). Additionally, engagement of the type II FcγR pathway by sFc has been shown to result in the up-regulation of FcγRIIB, release of the cytokine IL-33, and the expansion of T-regulatory cells (*3, 6, 8, 32*). More recently, sialylated IgG was shown to repress nuclear-factor kB (NF-kB) driven responses in influenza infection models through the induction of the transcription factor repressor element-1 silencing transcription factor (REST), protecting mice from severe lung inflammation, in a mechanism also dependent on the type II FcγR SIGN-R1 (*33*). However, the role of type II FcγRs in mediating anti-inflammatory activity has been challenged by other groups, with some arguing that DC-SIGN does not interact with human IgG (*34*) or sialylated glycans (*35*).

To characterize the requirement of type II FcγRs in the anti-inflammatory effect of V11 sFc, mice were administered V11 sFc prior to K/BxN serum injection, and mice received daily injections of a SIGN-R1 blocking antibody (*36*) or an isotype control (**Fig. 4A)**. Blocking SIGN-R1 resulted in the loss of anti-inflammatory activity of V11 sFc, while isotype control treated mice were protected from severe inflammation (**Fig. 4B-C)**, demonstrating that V11 sFc resolves inflammation in a type II FcγR dependent mechanism similar to IVIG (*9*). Recently, the C-type lectin Dectin-1 was shown to interact with FcγRIIB to stabilize its cell-surface expression and enhance ability to bind IVIG (*21*), suggesting C-type lectins such as type II FcγRs may directly interact with type I FcγRs on the cell surface. Interestingly, we found that co-expressing DC-SIGN and FcγRIIB in HEK 293-T cells enhance the expression level of FcγRIIB on the cell surface (**Fig. 4D, fig. S7A-B)**, indicating DC-SIGN may directly interact with FcγRIIB. Type I FcγRs like FcγRIIB have multiple N-linked glycans on their ectodomains, consisting primarily of complex type glycans (*37-39*), which may potentially interact with DC-SIGN. We generated a full-length variant of FcγRIIB lacking these core glycosylation sites (FcγRIIB^Δ^) (N106Q, N180Q, N187Q) (**fig. S7C)** and performed co-transfection experiments with DC-SIGN. Knocking out glycans on FcγRIIB resulted in the loss of enhanced cell surface expression when co-expressed with DC-SIGN, suggesting DC-SIGN may be binding N-linked glycans on FcγRIIB and stabilizing its expression on the cell surface. We found that another major type II FcγR, CD23 (FcεRII), also enhances the cell surface expression of FcγRIIB, when co-expressed, and this enhancement is dependent upon the glycans on FcγRIIB as well (**fig. S7F-H)**.

**Figure 4:**
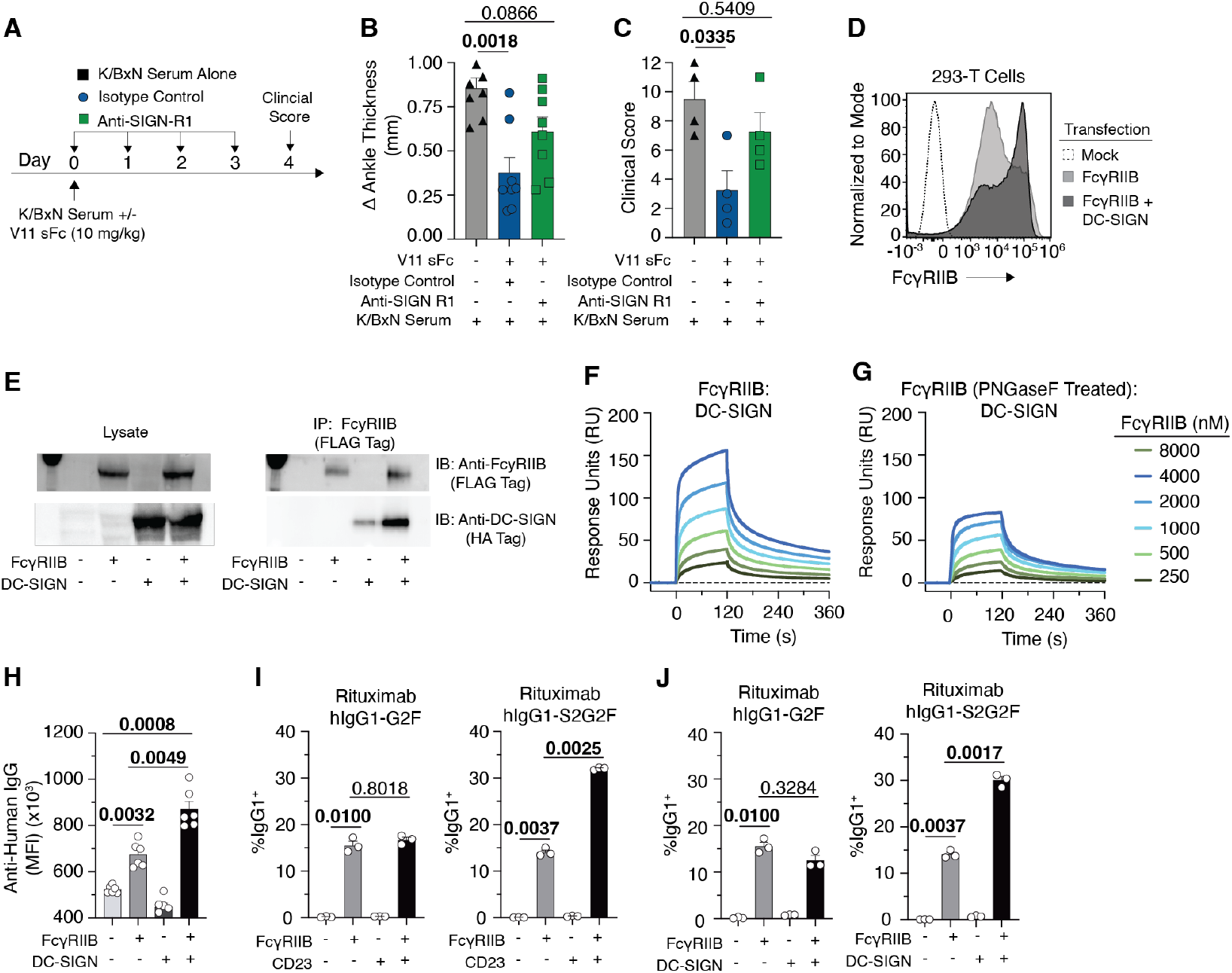
Type II FcγRs are required for V11 sFc mediated anti-inflammatory activity and synergize with type I FcγRs to bind IgG. **(A)** hFcγR mice (4/group) were dosed with 10 mg/kg of V11 sFc before K/BxN serum injection. Mice were then treated daily with an isotype control or a SIGN-R1 blocking antibody. Change in ankle thickness (**B**) or clinical score **(C)** four days post K/BxN serum injection. **(D)** Histogram of FcγRIIB expression in HEK 293-T cells transfected with FcγRIIB alone or in combination with DC-SIGN. **(E)** Western blot analysis of cell lysates or anti-FLAG tag immunoprecipitants of 293-F cells transfected with FLAG tagged FcγRIIB and/or HA-tagged DC-SIGN. Immunoblot (IB). (**F-G)** SPR binding of serial dilutions of FcγRIIB or PNGaseF treated FcγRIIB reacting against immobilized DC-SIGN. **(H)** Binding of fluorescently labeled IVIG (100 ug/ml) incubated with FcγRIIB and/or DC-SIGN expressing cells, measured by flow cytometry. (**I-J**) Binding of Rituximab-G2F or -S2G2F to cells expressing FcγRIIB alone or in combination with CD23 or DC-SIGN. Brown-Forsythe and Welch ANOVA tests, followed by Dunnett’s T3 multiple comparisons test was performed for (B), (C), (H), (I), and (J). P-values indicated above groups, significant values in bold.

To determine if DC-SIGN physically interacts with FcγRIIB, we next performed co-immunoprecipitation experiments in which FLAG-tagged FcγRIIB and HA-tagged DC-SIGN were expressed alone or in combination in HEK 293 cells. One day post transfection, cells were lysed and FcγRIIB was immunoprecipitated (IP) with anti-FLAG tag beads. Cell lysates and IP elutions were blotted to detect FcγRIIB (anti-FLAG tag) or DC-SIGN (anti-HA). We found similar levels of FcγRIIB and DC-SIGN expression in single expressing or co-expressing cells (**Fig. 4E)**. DC-SIGN co-immunoprecipitated with FcγRIIB in cells expressing both receptors indicating a direct physical interaction between DC-SIGN and FcγRIIB, although non-specific binding of DC-SIGN itself to anti-FLAG beads was observed as well. To further characterize a physical interaction between DC-SIGN and FcγRIIB, we reacted recombinant FcγRIIB ectodomains against DC-SIGN covalently coupled to a SPR chip. FcγRIIB directly interacted with immobilized DC-SIGN, and this interaction was markedly reduced when treating FcγRIIB with PNGaseF which cleaves N-linked glycans from proteins (**Fig. 4F-G)**, suggesting DC-SIGN may interact with both glycans and the protein backbone of FcγRIIB.

As the cell surface expression of FcγRIIB is enhanced upon co-expression with DC-SIGN, we next determined if this enhanced expression leads to enhanced binding of IgG. Fluorescently labeled IVIG was incubated with FcγRIIB and or DC-SIGN expressing cells and binding was measured by flow cytometry. As expected, FcγRIIB^+^DC-SIGN^+^ cells have an enhanced ability to bind fluorescently labeled monomeric IVIG compared to FcγRIIB^+^ only expressing cells (**Fig. 4H)**. Notably, DC-SIGN^+^ only expressing cells did not bind IVIG at this concentration, indicating that DC-SIGN functions to augment FcγRIIB binding to IgG when both are expressed together.

Next, we fluorescently labeled the anti-CD20 monoclonal antibody Rituximab, glyco-engineered to be homogenously galactosylated (G2F) or sialylated (S2G2F) (*40*). We incubated labeled IgGs with cells expressing FcγRIIB alone or in combination with CD23 or DC-SIGN and found that only the sialylated form of Rituximab exhibited enhanced binding to type I and II FcγR co-expressing cells, suggesting that co-expression of type I and II FcγRs drives the binding of sialylated IgG (**Fig. 4I-J, fig. S7I-J)**.

Thus, these studies suggest a novel function of DC-SIGN, and by extension other type II FcγRs, in which type II FcγRs directly interact with type I FcγRs to augment their binding to IgG, and this interaction is critical for mediating the anti-inflammatory activity of IVIG or sialylated IgG. Our data further suggests that FcγRIIB contributes to the anti-inflammatory activity of sialylated IgG through its ITIM domain. FcγRIIA, an ITAM containing Type I FcγR, is 92% homologous to FcγRIIB in its extracellular domain and is capable of associating with DC-SIGN (**fig. S8**), similar to FcγRIIB. However, enhancing FcγRIIA binding by the GA Fc variant did not result in anti-inflammatory activity when this variant was sialylated (**Fig. 2B-C**), while enhancing FcγRIIB binding alone, even in the absence of sialylation, resulted in a partial anti-inflammatory phenotype **(fig. S6)**. This finding may explain studies challenging the role in DC-SIGN interacting with sialylated IgG or IVIG, as these studies only characterized the binding of IgG by DC-SIGN alone or type I FcγR expressing cells, not in combination (*34*), as type II FcγRs are expressed on cells which also express type I FcγRs, and these experiments suggest type I and II FcγRs are found clustered together on the cell surface and signal IgG (*11, 21*).

Our findings demonstrate that sialylated IgG engineered to have enhanced affinity to the inhibitory FcγRIIB has potent *in vivo* anti-inflammatory activity, effective at a dose at least 100-fold lower than conventional IVIG therapy in multiple mouse models of autoimmunity.

Additionally, we show that this activity is dependent on the type II FcγR DC-SIGN and its mouse ortholog SIGN-R1 and describe a new function of type II FcγRs, which is to directly interact with type I FcγRs, augmenting their ability to bind and signal sialylated IgG. We propose a model in which V11 sFc engages in similar pathways as IVIG and sFc mediating anti-inflammatory activity, and through enhancing the affinity sFc for FcγRIIB, the effective therapeutic dose of sFc can be reduced at least 100-fold as compared to IVIG (**fig. S9)** (*41*). Recombinantly produced V11 sFc thus represents an attractive replacement for IVIG therapy for a wide array of autoimmune disorders currently being treated with IVIG, clinically effective at substantially lower doses compared to IVIG. Its anti-inflammatory properties do not result in depletion of IgG, as seen with FcRn blockade or B cell depletion therapies, thereby retaining the protective, anti-microbial properties of serum IgG, while mitigating the hyper-inflammatory sequalae of autoimmune diseases.

## Supporting information

Jones et. al Supplement

## Acknowledgments

We thank E. Lam, B. Bhagwandin-Colosi, A. Mozqueda, and R. Peraza (Rockefeller University) for excellent technical support. We thank Lai-Xi Wang (University of Maryland) for providing glycoengineered Rituximab antibodies.

## Author Contributions

ATJ, SB, and JVR designed the study. ATJ, TM, and AEM performed experiments, collected, and analyzed data. ATJ, TM, and JVR wrote the manuscript.

## Data and Materials Availability

All data are available in the main text or the supplementary materials. All materials are available on request after completion of a materials transfer agreement with The Rockefeller University.

