## Supplementary material for "The anti-inflammatory activity of IgG requires the co-engagement of types I and II Fc receptors": Jones et. al Supplement

**The PDF file includes:**

Materials and Methods
Figs. S1 to S9

### Materials and Methods

#### Recombinant protein generation

Recombinant human IgG1 Fc proteins, consisting of the full length IgG1 hinge, CHγ2, and CHγ3 domains, were produced via transient transfection of HEK 293-F cells (ThermoFisher, Cat no: A14635) and purified from cell-fire supernatants by affinity chromatography using Protein G Sepharose beads (GE Healthcare). To remove aggregates, Fc proteins were subsequently re-purified and dialyzed into phosphate-buffered saline (PBS) via size exclusion chromatography using a HiLoad 16/600 Superdex 200 pg resin column on an AKTA Pure (Cytivia) chromatography system. Monomeric Fc proteins were then concentrated with Amicon Ultra Centrifugal Filters (Millipore), filter-sterilized (0.22 μm), and quantified via BCA protein assay (ThermoFisher).

Sialylated Fc proteins were generated by co-transfecting HEK 293-F cells with Fc expression vectors and plasmids encoding open reading frames of Human Beta-1,4-galactosyltransferase 1 (B4GALT1) (Invivogen, Cat no: puno1-hb4galt1), and Human Beta-galactoside alpha-2,6-sialyltransferase 1 (ST6GAL1) (Invivogen, Cat no: puno1-hst6gal1). The human Fc variants GRLR (G236R/L328R), GA (G236A), or V11 (G237D, P238D, H268D, P271G, A330R) were generated via site-directed mutagenesis or as synthetic genes (IDT), and expressed and purified similar to sialylated WT Fc proteins. Prior to *in vivo* experiments, the monomeric purity of the Fc proteins was reassessed via size exclusion chromatography with a Superdex 200 Increase 10/300 GL column. All Fc proteins were at least 95% monomeric protein. To generate asialylated Fc proteins, sialylated Fc proteins were treated with α2-3,6,8 Neuraminidase (NEB) overnight and re-purified via size exclusion chromatography.

Recombinant human FcγRs were generated as his-tagged ectodomains based derived from annotated (Uniprot) sequences, expressed as soluble proteins via transient transfection of 293-F cells, and purified by affinity chromatography using HisTrap HP resin columns, followed by size exclusion chromatography to isolate monomeric FcγRs.

#### Animal Studies

All in vivo experiments were performed in compliance with federal laws and institutional guidelines and have been approved by The Rockefeller University Institutional Animal Care and Use Committee (Protocol number 20029-H). Mice were bred and maintained in the Comparative Bioscience Center at The Rockefeller University. FcγR Humanized (hFcγR) mice (FcγR$\alpha_{null}$, hFcγRI$^{+}$, hFcγRIIAR$^{131+}$, hFcγRIIB$^{+}$, FcγRIIIAF$^{158+}$, FcγRIIIB$^{+}$) were generated on the C57BL/6 background and have been extensively characterized in previous studies (*12, 20*).

K/BxN serum transfer induced arthritis was conducted as previously described (*4*). Briefly, KRN TCR transgenic mice (a gift from D. Mathis and C. Benoist, Harvard Medical School, Boston Massachusetts, USA) were bred with male NOD mice (strain 032445, Jackson Laboratory) to generate K/BxN mice which spontaneously develop arthritis (*15*). Serum from arthritic mice was collected, pooled together, and frozen. To induce arthritis in naive, 100 μl K/BxN serum was injected intraperitoneally in seven-eight week old male hFcγR$^{+}$ mice, and the development of arthritis and inflammation was evaluated through clinical scoring and measurement of ankle thickness using a digital caliper. To characterize the prophylactic anti-inflammatory activity of

IVIG or Fc proteins, male hFcγR mice were injected intravenously with IVIG or Fc proteins at the indicated dose one hour prior to intraperitoneal injection with 100 μl of K/BxN serum. For clinical scoring, a score of 0-3 was given to each paw, a score of 0 being no inflammation and 3 representing severe swelling and inflammation. Scores from each paw were summed together to give an overall clinical score.

For SIGN-R1 blocking experiments, male hFcγR+ mice were treated with 10 mg/kg of V11 sFc intravenously one hour before intraperitoneal injection with 100 μl of K/BxN serum. To block SIGN-R1, mice were injected subcutaneously with 100 μg of an anti-SIGN-R1 antibody (clone 22D1, Bioxcell, Cat no: BE0220) 24 hours prior to sFc and K/BxN serum injection. Mice received two additional injections with anti-SIGN-R1 on days 1 and 3 post serum transfer. A control group of mice was injected with an isotype control (Bioxcell, Cat no: BE0091) at the same dose and schedule as anti-SIGN-R1 treatment.

Experimental autoimmune encephalomyelitis (EAE) was induced with 8-to-10 week old $hFc\gamma R^+$ mice by immunizing mice subcutaneously with 200 μl of an emulsion consisting of $MOG_{35-55}$ antigen (MEVGWYRSPFSRVVHLYRNGK, Hooke Labs) emulsified in complete Freund's adjuvant (Hooke Labs, Cat no: EK-2110). Mice were subsequently intraperitoneally injected with 100 ng of pertussis toxin (Hooke Labs) 4 and 24 hours post immunization. Development of disease was monitored daily according to the following criteria: 0, no clinical signs; 0.5, partial loss of tail tone; 1, paralyzed tail; 2, loss of coordinated movement, and hind limb paresis; 2.5 one hind limb completely paralyzed; 3, both hind limbs paralyzed; 3.5, hind limbs paralyzed and/or hunched back and disorientation; 4, severely hunched back and weakness in forelimbs; 4.5, forelimbs paralyzed; 5, moribund. To characterize the anti-inflammatory effect of V11 sFc in the EAE model, male $hFc\gamma R^+$ mice were subcutaneously injected with 10 mg/kg of V11 sFc on days 5, 10, 15, and 20 post immunization.

**Tissue imaging**

To characterize inflammation in joints of V11 sFc treated and untreated K/BxN serum induced arthritic mice, hind legs were collected from euthanized mice and fixed overnight in 10% Neutral Buffered Formalin (NBF). Hind legs were subsequently de-calcified for a week (Decal, Statlab) before embedding in Tissue-Tek O.C.T and snap freezing to -80$^o$C. Frozen ankles were sectioned (10 um) on a Cryostat (Leica), dried overnight, and stained with a hematoxylin and eosin stain kit (Vector Labs).

To collect brains and spinal cords in EAE experiments, mice were euthanized and perfused via intra-cardiac puncture with cold PBS prior to perfusion with 10% Neutral Buffered Formalin and central nervous system tissues were harvested and stored in 10% NBF overnight. Following cryoprotection by incubation in a series of 15% then 30% sucrose/PBS solution overnight at 4 °C, the brain and spinal cords were then embedded in Tissue-Tek O.C.T. compound (Sakura) and cut on a cryostat. Serial sections (10-20 um thickness, 7 series) were collected on positively charged microscope slides (VWR) dried overnight and stored at -80$^o$C until further processing.

For immunofluorescence imaging, spinal cord sections were stained with a Myelin Basic Protein (MBP) antibody (1:150) overnight before washing and staining with a fluorescent secondary antibody for one hour followed by mounting with ProLong Gold Antifade Mountant with DAPI (Invitrogen, P36931)

All sample acquisition and analysis were conducted using the AKOYA PhenoCycler®-Fusion 2.0 and QuPath v0.5.1.

**Western Blotting**

Recombinant Fc proteins (5 ug) were resolved on SDS-PAGE gels in non-reducing conditions, transferred to PVDF membranes (BioRad) and blocked for one hour with PBS + 1% BSA. Blots were then incubated overnight with biotinylated SNA lectin (2 ug/ml, Vector Labs, Cat no: B-1305-2) or ECL (5 ug/ml, Vector Labs, Cat no: B-1145-5), followed by incubation with Peroxidase Streptavidin (1:50,000, Jackson ImmunoResearch, Cat no: 016-030-084). Blots were developed with a chemiluminescent substrate (Clarity ECL, Bio-Rad) and imaged on a ChemiDoc MP imaging system (BioRad). Parallel to SNA and ECL blots, 5 ug of Fc proteins were resolved via non-reducing SDS-PAGE gels and stained with SimplyBlue SafeStain (ThermoFisher) to visualize total protein. Gels were imaged on a ChemiDoc MP system. Blots and protein gels were analyzed on ImageJ software. To estimate the degree of sialylation or galactosylation, SNA and ECL band intensities were normalized protein gel loading controls and plotted as relative intensities.

**Immunoprecipitation**

HEK 293-F cells were transiently transfected with FLAG-tagged FcγRIIB (Sino, Cat no: HG10260-NF), FLAG-tagged FcγRIIA, (Sino, HG10374-NF) and/or HA-tagged DC-SIGN (Sino, Cat no: HG10200-CY) full-length open reading frame expression vectors. Two days post transfection, cells were lysed with IP-lysis buffer (Pierce) with EDTA-free protease inhibitors added (Roche) and clarified cell lysates were immunoprecipitated (IP) using anti-FLAG tag magnetic agarose (ThermoFisher, Cat no: A36797). Cell lysates and IP elutions were resolved under reducing conditions on SDS-PAGE gels, transferred to PVDF membranes, and blocked for one hour with TBS + 1% BSA. Blots were subsequently incubated overnight with a fluorescently labeled anti-FLAG tag antibody (ThermoFisher, Cat no: MA1-91878-D650) or a biotinylated anti-HA tag antibody (ThermoFisher, Cat no: 26183-BTIN). FLAG-tag immunoblots were fluorescently imaged on a Chemidoc MP imaging system. HA-tag immunoblots were washed and incubated with Peroxidase Streptavidin (1:50,0000, Jackson ImmunoResearch) before chemiluminescent imaging.

**SPR**

All experiments were performed with a Biacore T200 SPR System (Cytivia) at 25$^{o}$C. Fc proteins were immobilized on Series S Protein G sensor chips (Cytivia) in HBS-EP$^{+}$ Buffer (10 mM HEPES, pH 7.4, 150 mM NaCl, 3.4 mM EDTA, 0.005% (v/v) surfactant P20). Serial dilutions of recombinant human FcγRs were injected to the flow cells at 30 ul min$^{-1}$, with concentrations ranging from 8000 nM to 125 nM (1:2 serial dilutions) or 300 nM to 4.69 nM (1:2 serial dilutions) for the high affinity FcγRIA. FcγRs were allowed to associate for 120 seconds with Protein-G immobilized Fc proteins before a 900 second dissociation step. At the end of each cycle, the sensor chip was regenerated with glycine HCL buffer (10 mM, pH 1.5, 50 ul min$^{-1}$, 30 seconds). Background binding to blank immobilized flow cells was subtracted and affinity constants (steady-state affinity, or 1:1 fit for FcγRIA) was calculated using BIAcore T200 evaluation software (Cytivia).

Recombinant DC-SIGN (AcroBiosystems, Cat no: CD9-H5246-100ug) was immobilized to a Series S CM5 chip (Cytivia) using an amine couple kit (Cytivia) in HBS-$P^+$ Buffer (10 mM HEPES, pH 7.4, 150 mM NaCl, 0.005% (v/v) surfactant P20) supplemented with 10 mM $CaCl_2$ and 10 mM $MgCl_2$. Serial dilutions of recombinant native or PNGaseF treated (NEB, Cat no: P0706S) human FcγRIIA$^{R131}$ or FcγRIIB were injected into flow cells and associated against immobilized DC-SIGN for 120 seconds, followed by a 900 second dissociation step, and subsequent regeneration with 10 mM glycine HCL buffer (pH 1.5). Background binding to non-immobilized flow cells was subtracted and affinity constants were calculated using BIAcore T200 evaluation software (1:1 fit).

**Flow cytometry**

Expi293 cell lines transiently transfected to express full length FcγRIIB, FcγRIIA$^{H131}$, human CD23, or human DC-SIGN alone or in combination were harvested and stained with fluorescently labeled anti-human FcγRIIB (1 uglml, clone 2B6), anti-human FcγRIIA (1 ug/ml clone IV.3), anti-DC-SIGN (1:200, BD, clone DCN46), anti-human CD23 (1:200, BD, clone M-L233), and a fixable live/dead stain (ThermoFisher). Staining was performed at 4$^{o}$C for 30 minutes. Cells were washed twice and analyzed on an Attune NxT flow cytometry (Thermofisher) using Attune NxT software and data was analyzed using FlowJo software.

For IgG binding studies, IVIG or glycoengineered Rituximab G2F or S2G2F (*40*) were fluorescently labeled with Dylight antibody labeling kits (ThermoFisher) and incubated overnight at 4$^{o}$C (10 ug/ml) with transfected FcγR expressing 293 cell lines in 10 mM Hepes, 150 mM NaCl, 10 mM $CaCl_2$, 10 mM $MgCl_2$, 0.5% BSA, pH 7.4 buffer. Cells were immediately fixed in 2% paraformaldehyde for 20 minutes at 4$^{o}$C before washing and staining with fluorescent anti-FcγR antibodies and a fixable live/dead stain. Cells were subsequently washed twice and analyzed on an Attune NxT flow cytometer as before, and IgG binding was determined through gating on live FcγR+ cells and measuring the median fluorescent intensity (MFI) (IVIG) or percent IgG1+ cells (Rituximab G2F and S2G2F IgG1).

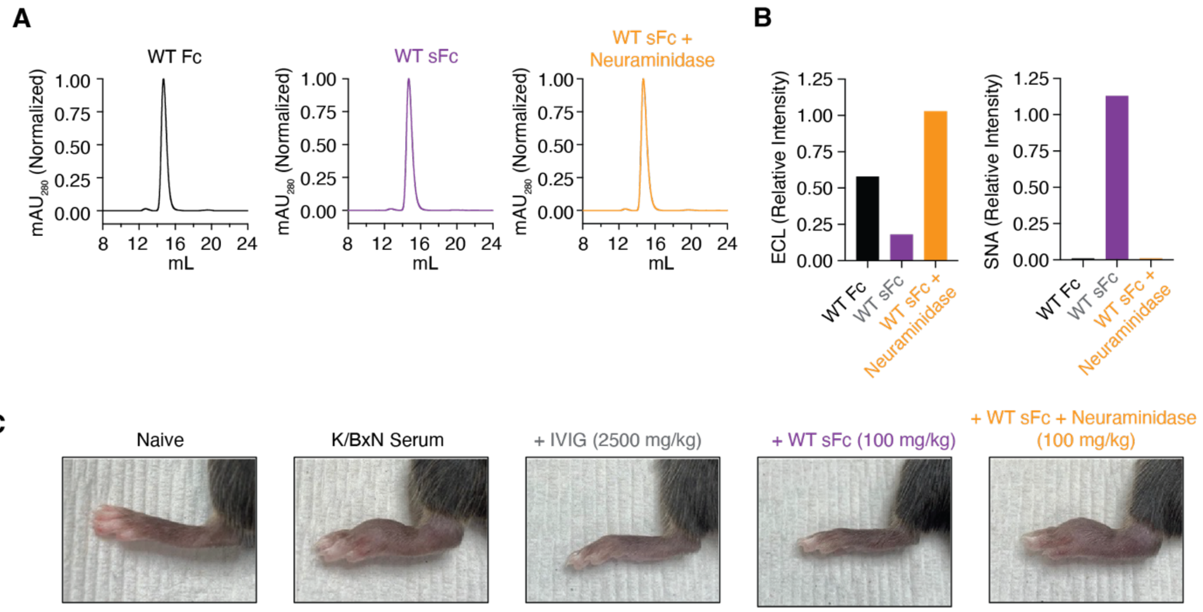


**Supplemental Figure 1: (A).** Size exclusion chromatography (SEC) analysis of WT Fc, WT sFc, and WT sFc + Neuraminidase Fc proteins. **(B)** Relative intensity in ECL and SNA lectin blots in (Fig 1B). **(C)** Images of ankles in mice from study in (Fig 1C), taken four days post K/BxN serum injection.

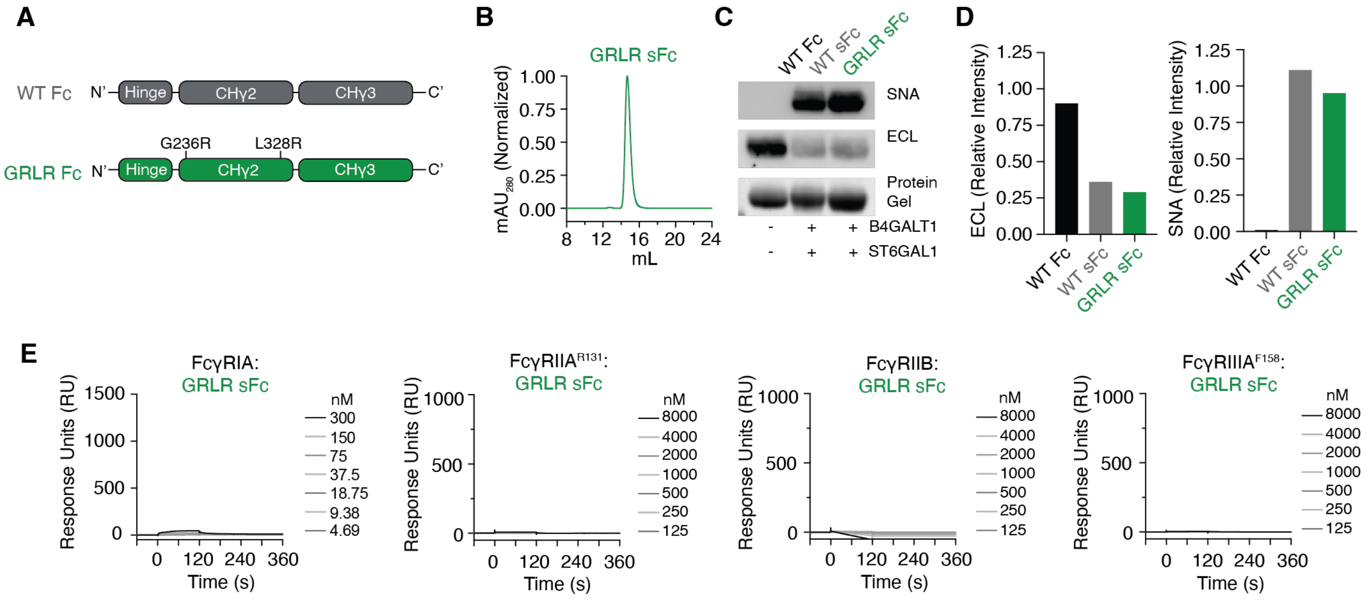


**Supplemental Figure 2: (A).** Schematic of WT Fc and GRLR (G236R, L328R) Fc expression vectors. **(B)** SEC analysis of GRLR sFc proteins. **(C)** Lectin blot analysis of WT Fc, WT sFc, and GRLR sFc proteins with ECL and SNA. **(D)** Relative intensities of ECL and SNA lectin blots in (C). **(E).** SPR analysis of protein-G immobilized GRLR sFc reacted with type I FcγRs.

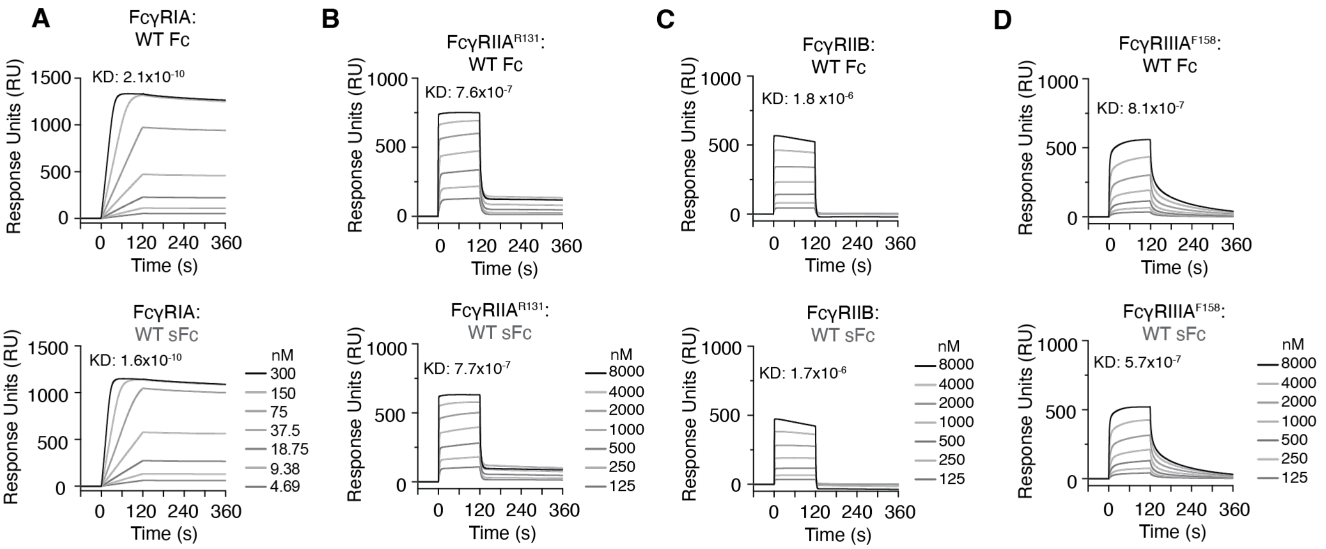


**Supplemental Figure 3:** Surface plasmon resonance (SPR) of protein-G immobilized WT Fc and WT sFc proteins reacted against **(A)** FcγRIA, **(B)** FcγRIIA$^{R131}$, **(C)** FcγRIIB, **(D)** FcγRIIIA$^{F158}$.

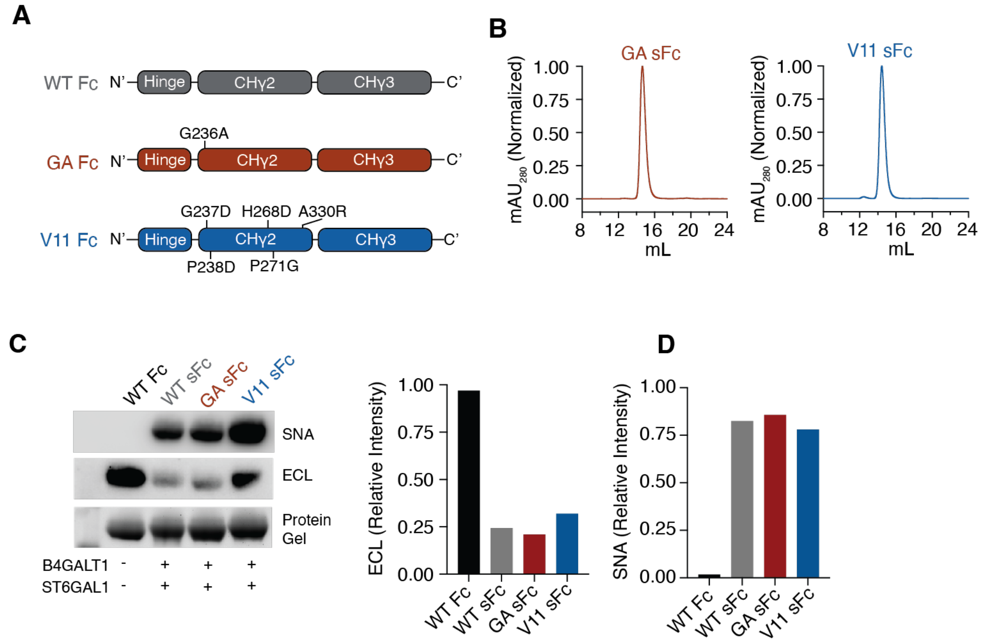


**Supplemental Figure 4: (A)** Schematic of WT Fc, GA (G236A) Fc, or V11 Fc (G237D, P238D, H268D, P271G, A330R) expression vectors. **(B)** SEC analysis of GA sFc and V11 sFc proteins. **(C)** SNA and ECL lectin western blots and protein gel for WT Fc, WT sFc, GA sFc, and V11 sFc proteins. **(D)** Relative intensities of ECL and SNA blots in **(C)**.

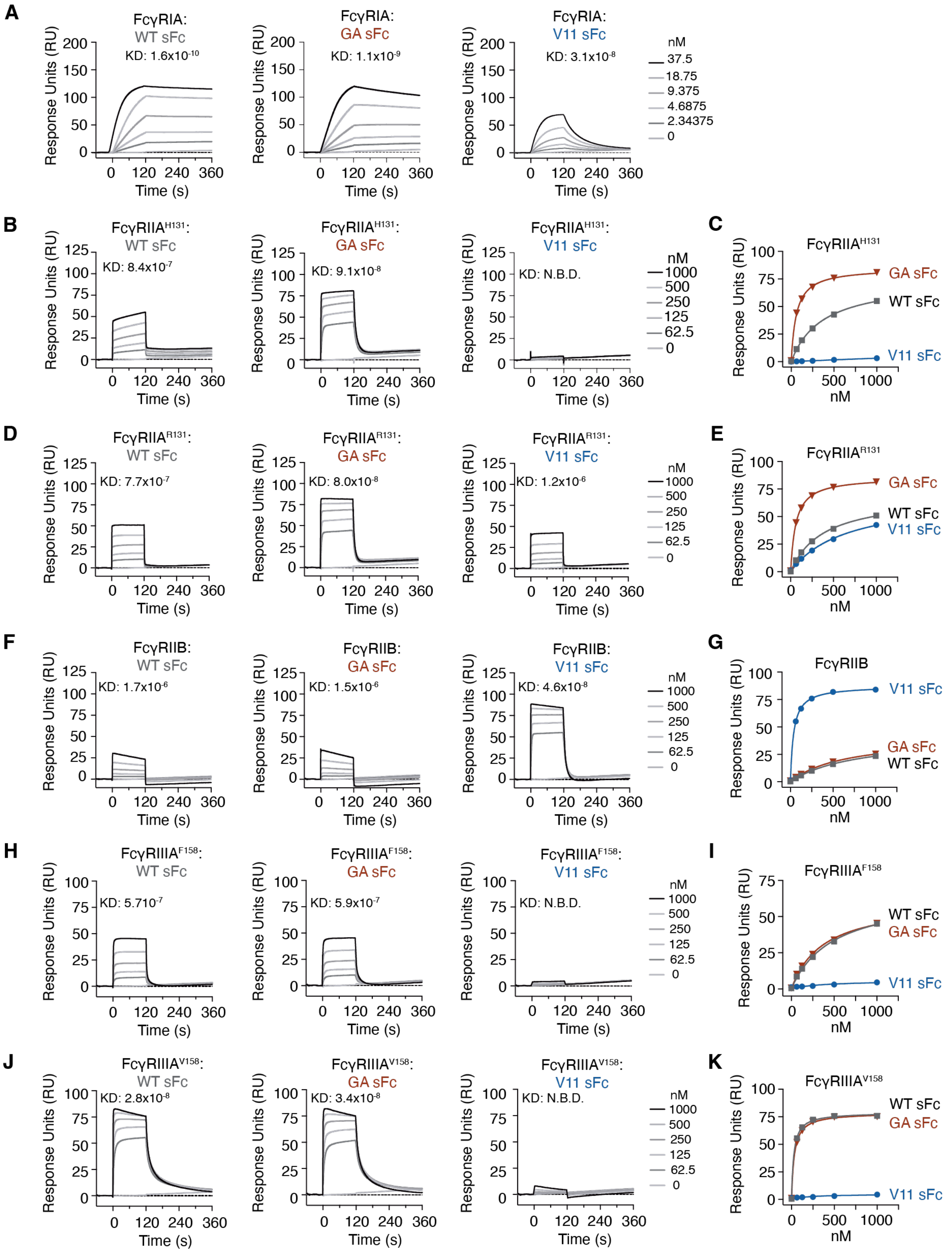


**Supplemental Figure 5:** SPR analysis Protein-G immobilized WT sFc, GA sFc, or V11 sFc reacted with **(A)** FcγRIA, **(B)** FcγRIIA$^{H131}$, **(D)** FcγRIIA$^{R131}$, **(F)** FcγRIIB, **(H)** FcγRIIIA$^{F158}$, and **(J)** FcγRIIIA$^{V158}$. Steady-state affinity plots for representative FcγRs plotted in **(C)**, **(E)**, **(G)**, **(I)**, and **(K)**.

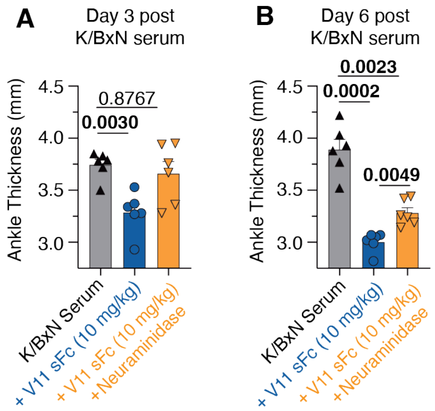


**Supplementary Figure 6:** Mice were dosed with 10 mg/kg of V11 sFc or neuraminidase treated V11 sFc one hour prior to K/BxN serum injection. Ankles were measured three days (A) and six days (B) post K/BxN serum induced arthritis. Brown-Forsythe and Welch ANOVA tests, followed by Dunnett's T3 multiple comparisons test was performed for **(A)** and **(B).**

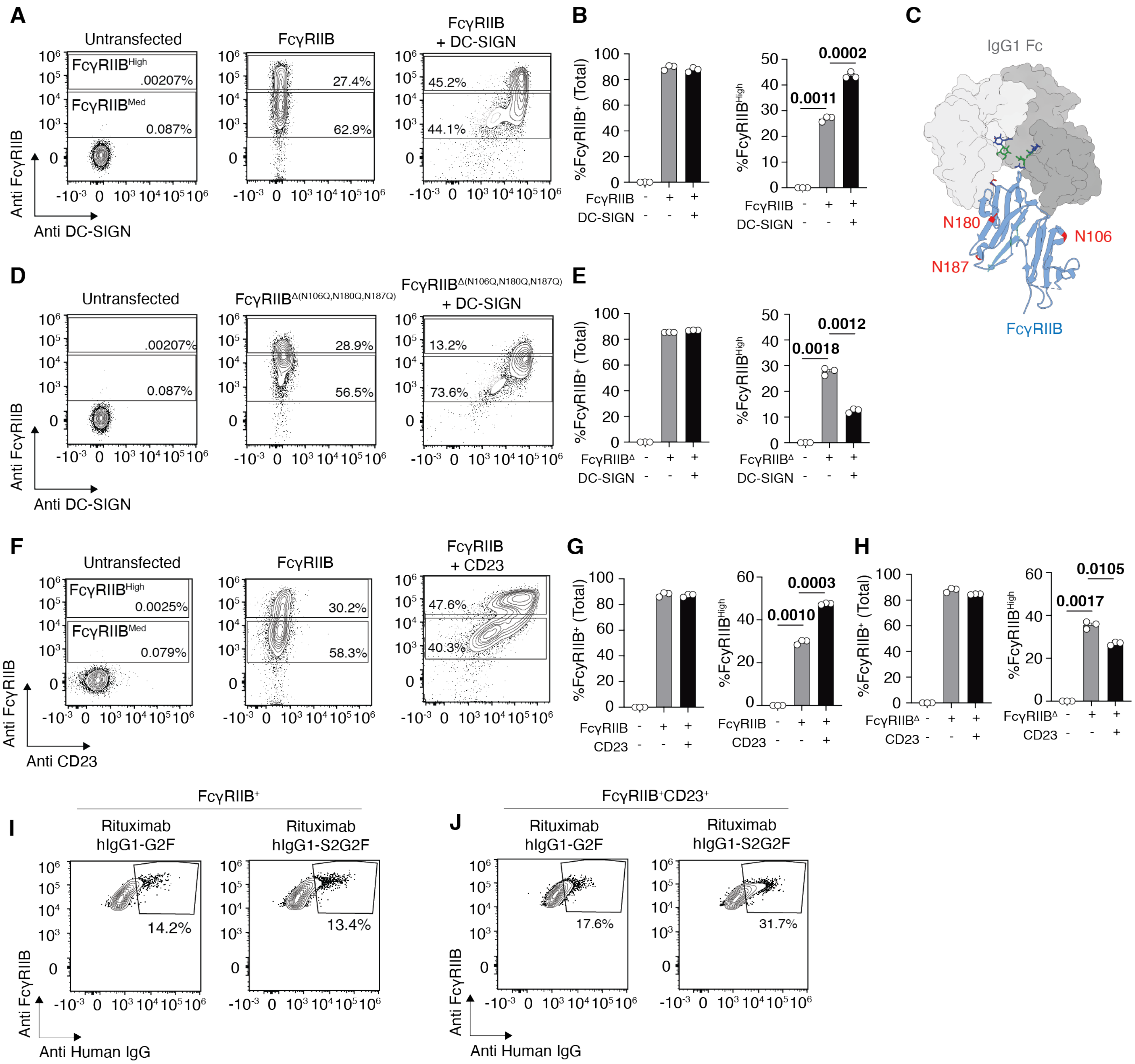


**Supplementary Figure 7:** Expi293 cells were mock transfected or transfected with full-length FcγRIIB alone or in combination with full-length DC-SIGN. Expression of FcγRIIB was measured by flow cytometry two days post transfection. **(A)** Gating strategy of FcγRIIB$^{med}$ and FcγRIIB$^{high}$ cells. Cells first first gated on live singlets. **(B)** Frequencies of % FcγRIIB$^{med}$ and %FcγRIIB$^{high}$ in transfected cells. **(C)** Structure of FcγRIIB in complex with IgG1 Fc (PBD 3WJJ). N-glycan sites on FcγRIIB are highlighted in red. **(D-E).** Expression of glycan knocked-out FcγRIIB (FcγRIIB$^{\Delta}$, N106Q, N180Q, N187Q) transfected alone or in combination with DC-SIGN. **(F-G).** Expression pattern of FcγRIIB or FcγRIIB$^{\Delta}$, expressed alone or in combination with Human CD23. Brown-Forsythe and Welch ANOVA tests, followed by Dunnett's T3 multiple comparisons test. **(I-J)**. Gating strategy of fluorescently labeled Rituximab G2F or S2G2F binding to FcγRIIB$^{+}$ or FcγRIIB$^{+}$CD23$^{+}$ expressing cells.

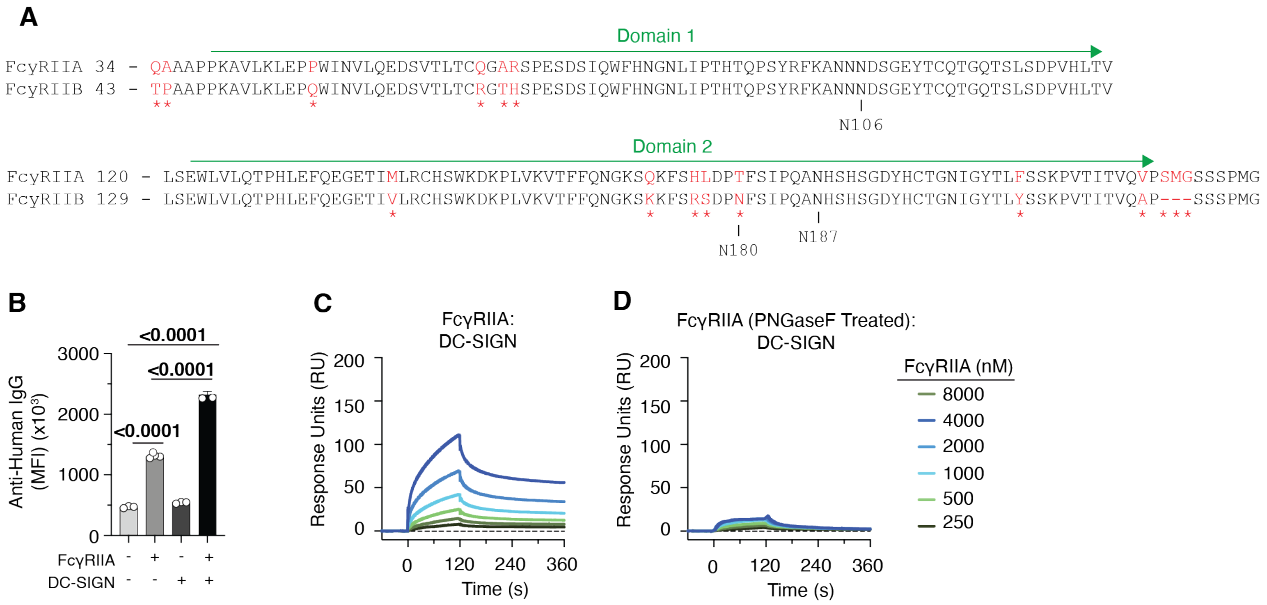


**Supplementary Figure 8: (A)** Protein alignment of ectodomains of FcγRIIA and FcγRIIB. Locations of N-glycan sites on FcγRIIB are indicated. **(B).** Binding of fluorescently labeled IVIG to cells transfected with FcγRIIA and DC-SIGN alone or in combination. **(C-D)** SPR of native or PNGaseF treated FcγRIIA reacting to immobilized human DC-SIGN.

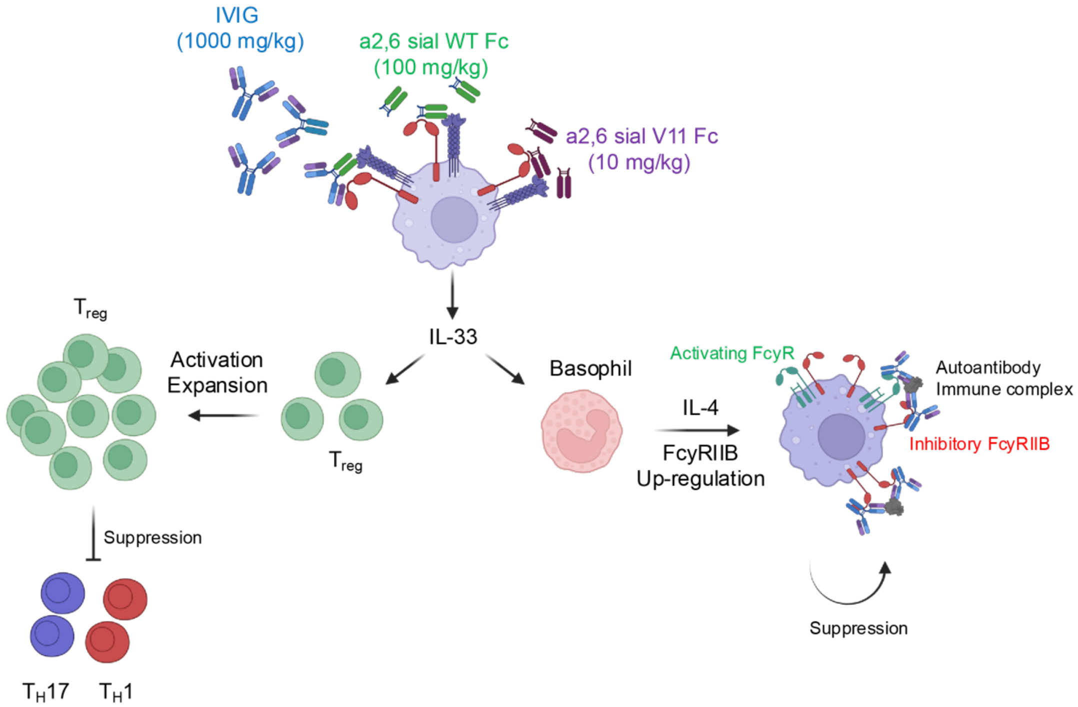


**Supplementary Figure 9:** Proposed mechanism of sialylated IgG mediated anti-inflammatory activity. IVIG (1000 mg/kg), WT sFc (100 mg/kg), or V11 sFc (10 mg/kg) engages the FcγRIIB pathway, leading to the release of the $T_H2$ cytokine IL-33, which acts in promoting the expansion of Treg cells or the release of IL-4 by basophils. IL-4 release results in the up-regulation of FcγRIIB on effector innate myeloid cells, increasing their threshold for FcγR-mediated cell activation.
